# Development of a non-invasive method for testicular toxicity evaluation using a novel compact magnetic resonance imaging system

**DOI:** 10.1101/2022.08.17.504248

**Authors:** Satoshi Yokota, Hidenobu Miyaso, Toshinori Hirai, Kousuke Suga, Tomohiko Wakayama, Yuhji Taquahashi, Satoshi Kitajima

**Affiliations:** Division of Cellular & Molecular Toxicology, Center for Biological Safety & Research, National Institute of Health Sciences, 3-25-26 Tono-machi, Kawasaki-ku, Kawasaki, Kanagawa 210-9501, Japan; Department of Anatomy, School of Medicine, International University of Health and Welfare, 4-3 Kozunomori, Narita, Chiba 286-8686, Japan; Department of Diagnostic Radiology, Graduate School of Medical Sciences, Kumamoto University, 1-1-1 Honjo, Chuo-ku, Kumamoto 860-8556, Japan; Department of Histology, Graduate School of Medical Sciences, Kumamoto University, 1-1-1 Honjo, Chuo-ku, Kumamoto 860-8556, Japan

**Keywords:** magnetic resonance in vivo imaging, mice, busulfan, testicular toxicity

## Abstract

In non-clinical animal studies for drug discovery, histopathological evaluation is the most powerful tool to assess testicular toxicity. However, histological analysis is extremely invasive; many experimental animals are needed to evaluate changes in the pathology and anatomy of the testes over time. As an alternative, small animal magnetic resonance imaging (MRI) offers a non-invasive methodology to examine testicular toxicity without radiation. The present study demonstrated the suitability of a new, ready-to-use compact MRI platform using a high-field permanent magnet to assist with the evaluation of testicular toxicity. To validate the utility of the MRI platform, male mice were treated with busulfan (40 mg/kg, intraperitoneal injection). Tenty-eight days after treatment, both testes in busulfan-treated and control mice (n = 3/group) were non-invasively scanned in situ by MRI at 1 tesla. On a T1-weighted, 3D gradient-echo MRI sequences (voxel size: 0.23 × 0.23 × 0.50 mm), the total testicular volume in busulfan-treated mice was significantly smaller than in controls. On T1-weighted images, the signal intensity of the testes was significantly higher in busulfan-treated mice than in controls. The mice were sacrificed, and the testes were isolated for histopathological analysis. The weight of the testes in busulfan-treated mice significantly decreased, similar to the results of the non-invasive analysis. Additionally, periodic acid-Schiff stain–positive effusions were observed in the interstitium of the busulfan-treated mouse testes, potentially explaining T1 shortening due to a high concentration of glycoproteinaceous content. The present data demonstrated a rapid evaluation of testicular toxicity in vivo by compact MRI.

## Introduction

In the drug discovery process, a pipeline compound that can affect the size and weight of the testes points to a sign of serious testicular toxicity. Therefore, testicular toxicity must be adequately evaluated before the first clinical trial on humans. In testing male reproductive toxicity, using experimental animals is preferable to detect similar changes occurring in humans exposed to the same chemicals. Among all adverse findings, histopathology and the weight of a male reproductive organ are the best non-clinical endpoints to detect reproductive toxicity (Scialli et al., 2018, Sotos & Tokar, 2012). However, it is difficult to determine testicular size from the body surface. Furthermore, these data can only be collected by sacrificing a large number of animals. In addition, histological findings are generally not available in humans. To address these concerns, it is important to develop a non-invasive method to evaluate testicular toxicity in vivo.

Owing to its lack of ionizing radiation and high-contrast resolution, magnetic resonance imaging (MRI) with a high-frequency magnetic field is a powerful, non-invasive tool to detect and monitor the status of lesions with high sensitivity (Pichler et al., 2010). MRI can be applied to small animals in non-clinical toxicological studies. This approach enabled us to evaluate time-course changes in pathological and anatomical findings and their reversibility in the same animals. It can also detect the presence and location of induced lesions and/or deposition of abnormal proteins, assisting with the selection of sections for subsequent histopathological examination. In fact, MRI has been used to detect amyloid plaques by using amyloid-beta precursor protein transgenic mice (Zhang et al., 2004); this demonstrates the usefulness of MRI as a complementary tool for conventional histopathology. Thus, MRI widens the range of potential non-invasive imaging modalities, expanding the scope of non-clinical studies. However, the widespread use of MRI systems in non-clinical studies are hindered by obstacles, such as high costs of purchase and maintenance, significant siting and installation requirements, and complicated operations.

With progress in design technology, a novel compact MRI system with a high-field permanent magnet (∼1.0 tesla) has been developed to reduce the cost and complexity of superconducting magnets for MRI systems (Taketa et al., 2015, Pittala et al., 2018, Daneshgaran et al., 2019). The new system is portable and self-shielded; it can be placed in most institutions. Cryogens, plumbing, chemicals, and supplemental power suppliers or coolers are not required. Compared with conventional MRI systems (Natt et al., 2002), the new system has been utilized in several non-clinical studies, including those on hepato-, nephro-, and neurotoxicities (Taketa et al., 2015, Tempel-Brami et al., 2015). However, no evidence has been found on the evaluation of testicular toxicity in vivo by MRI.

The aim of the present study was to examine the usefulness of a ready-to-use, novel compact MRI platform in evaluating testicular toxicity. We used an infertility mouse model commonly used for busulfan administration (Simmons et al., 1975, Nakata et al., 2020, Gutierrez et al., 2016, Wang et al., 2010). Changes in the T1-weighted images of the testes were examined using a compact MRI system and compared with testicular histology results.

## Materials and Methods

### Animals

A total of six male C57BL/6J mice (CLEA Japan, Tokyo, Japan) aged 5 weeks were purchased and maintained under standard laboratory conditions. The mice were housed in individually ventilated cage systems (Lab Product Inc., Seaford, DE, USA) at 23 ± 2 °C, under a 12-hr light/dark cycle (lights on from 8:00 to 20:00), and with food and water provided ad libitum. Body weight was measured once a week (three mice per cage group). We followed the guidelines for animal care and use established by the Ethical Committee for Animal Experiments of the National Institutes of Health Sciences. The animal facility was approved by the Health Science Center for Accreditation of Laboratory Animal Care, Japan. All experimental protocols in the study were reviewed and approved by the Committee for Proper Experimental Animal Use and Welfare, a peer-review panel established at the NIHS (experimental approval no. 816).

### Experimental design

After 1 week of habituation, the mice were randomly divided into two groups: the vehicle control (n = 3) and busulfan-treated groups (n = 3). The vehicle control group received two intraperitoneal injections (i.p.) of a vehicle containing 10 % dimethyl sulfoxide (DMSO; 031-24051; CultureSure^®^; FUJIFILM Wako Pure Chemical Industries, Ltd., Osaka, Japan) in saline (Otsuka Pharmaceutical Co., Tokyo, Japan) with a 3-hr interval between injections. Busulfan (B-2635; Sigma–Aldrich, St. Louis, MO, USA) was dissolved in DMSO at a concentration of 10 mg/mL and was gradually added to nine times the volume of saline (final concentration: 1 mg/mL). As previously described, the busulfan-treated group received two i.p. (20 mL/kg per injection) of busulfan at a total dose of 40 mg/kg body weight (Xie et al., 2020). After 28 days, the mice were anesthetized with isoflurane for MRI analysis. After MRI data were obtained, the reproductive organs of male mice were collected under isoflurane anesthesia. All efforts were made to minimize animal suffering. Each mouse was weighed; blood was collected from the inferior vena cava, and the testes were removed. The dissected tissues were rapidly fixed for histopathological analysis.

### In vivo imaging and analysis using a compact MRI

Both busulfan-treated and control mice (n = 3/group) were anesthetized with isoflurane, and the testes were scanned using a compact MRI system (M7 Permanent Magnet System; 1.05 tesla; Aspect Imaging, Shoham, Israel) equipped with an application-specific mouse body radiofrequency coil. For in vivo imaging, the mice were maintained in an anesthetized state with 1.5% isoflurane in O_2_ and placed on a specially designed heated bed, where the respiratory rate was monitored to determine the anesthetic level (PC_SAM; SA Instruments, NY, USA).

High-resolution datasets were obtained using a T1-weighted, 3D gradient-echo MRI sequence (echo time/repetition time, 2.0 msec/20.0 msec; field of view, 27.04 × 27.04 mm; matrix, 117 × 117; voxel size, 0.23 × 0.23 × 0.50 mm; scan time, 119 sec). For quantitative evaluation, the signal-to-noise ratios of selected testis structures were determined. This was defined as the MRI signal intensity divided by the standard deviation of the noise. Testicular volume estimates were obtained by drawing 3D regions of interest in a sophisticated image processing and analysis software package (VivoQuant™ 2020; Invicro, Boston, MA, USA) that is fully integrated into the M7 imaging system.

### Testicular histology

For histopathological analysis, the testes were removed and immersed in Bouin’s fixative (023-17361; FUJIFILM Wako Pure Chemical Industries, Ltd., Osaka, Japan) for 48 hr. After fixation, the testes were processed in graded alcohol, cleared in xylene, and embedded in paraffin. Sections (4-**μ**m thick) were prepared, deparaffinized, and stained with periodic acid–Schiff–hematoxylin (PAS-H) to distinguish germ cell types and seminiferous tubule stages. The slides were mounted using Entellan® (Nacalai Tesque Inc., Kyoto, Japan) and air-dried prior to microscopic examination. A total of 100 seminiferous tubule sections were observed in two tissue sections, which were obtained from two discrete portions of a testis from a single mouse. We evaluated two tissue sections per mouse (n = 3 per group) by using a light microscope (BX50; Olympus Co., Tokyo, Japan) with the cellSens imaging software. The percentage of germ cell depletion in the seminiferous tubules was assessed.

### Statistical analysis

Data are presented as mean ± standard error of the mean (SEM). Welch’s t-test was used to detect significant differences between the control and busulfan-treated groups. Statistical significance was set at P <0.05.

## Results

### Body and organ weight

Busulfan treatment decreased body weight (control: 26.0 ± 0.4 g, busulfan-treated: 24.0 ± 0.2 g, P <0.05). Fig 1A and B shows the gross appearance of the testes in both control and busulfan-treated mice. The absolute weight of the testes in busulfan-treated mice was significantly lower than that in control mice (Fig. 1E).

**Figure 1.**
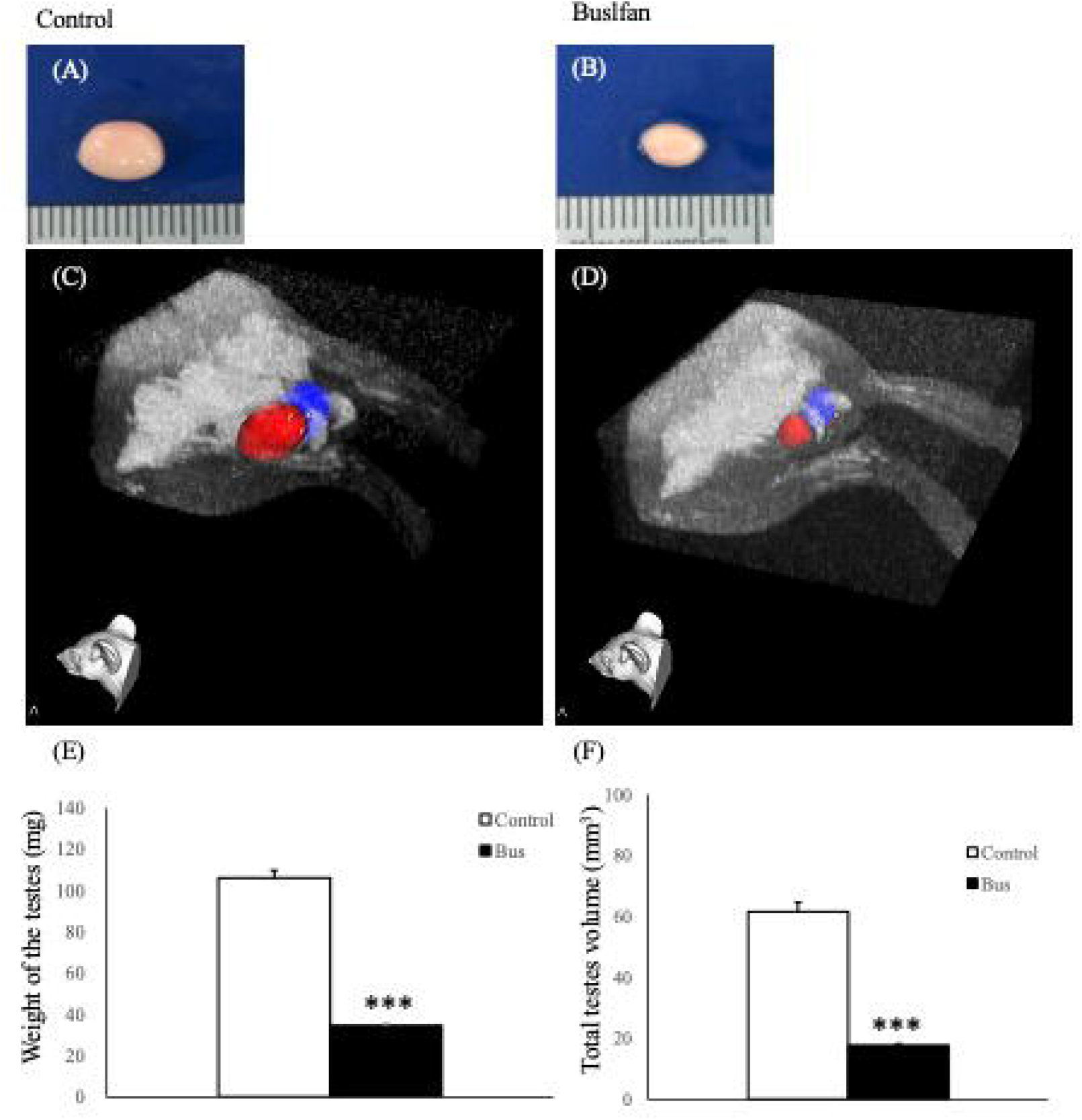
Photomicrographs and T1-weighted, 3D gradient-echo MR images of the testes. Photomicrographs of the gross appearances of (A) control and (B) busulfan-treated mouse testes. Coronal, maximum-intensity projection images of (C) control and (D) busulfan-treated mouse testes. High-resolution datasets were obtained with the following parameters: echo time/repetition time, 2.0 msec/20.0 msec; field of view, 27.04 × 27.04 mm; matrix, 117 × 117; voxel size, 0.23 × 0.23 × 0.50 mm; scan time, 119 sec. (E, F) Busulfan significantly decreased the weight and total volume of the testes. Data are expressed as mean ± SEM (n = 3 each). *P <0.05, **P <0.01, ***P <0.001 vs. control (Welch’s t-test).

### MRI analysis

Fig. 1C and D shows the maximum-intensity projection of T1-weighted, gradient-echo images of control and busulfan-treated mice, respectively. Measurements in the images showed a total testicular volume of 61.5 ± 3.2 mm^3^ (mean ± SEM) in the control mice and 17.6 ± 0.7 mm^3^ (mean ± SEM) in the busulfan-treated mice (Fig. 1F). Fig. 2A-D shows the T1-weighted images of mouse testes from which the signal was measured. The signal on T1-weighted images was significantly higher in busulfan-treated mice than in control mice. (Fig. 2E).

**Figure.**
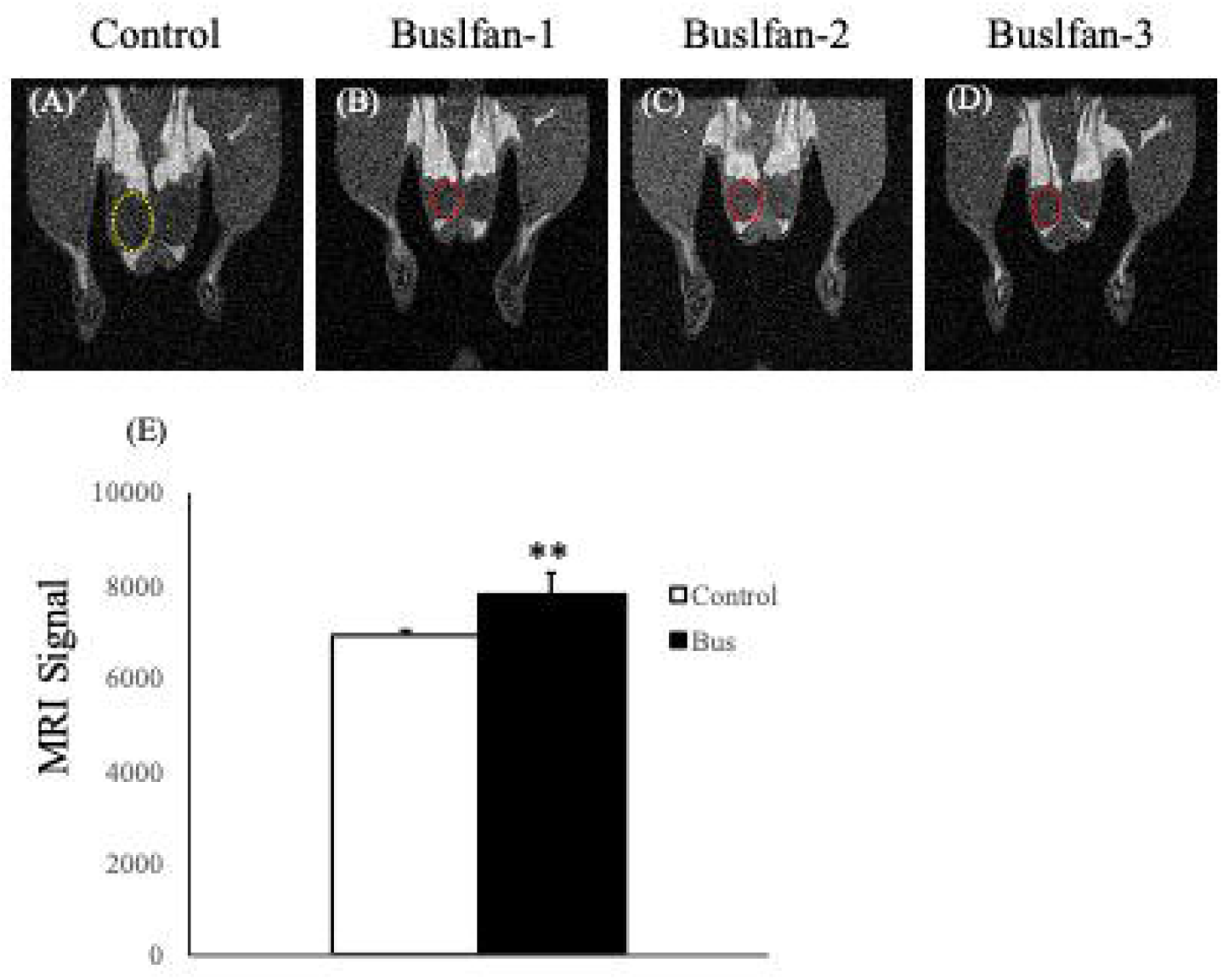
T1-weighted, 2D gradient-echo images of both control and busulfan-treated mice. T1-weighted, 2D gradient-echo images showing the regions of interest in the testes of (A) control mice, indicated by the yellow oval, (B-D) and busulfan-treated mice, indicated by the red ovals. (E) Compared to that of control mice, the signal of the testes on T1-weighted images significantly increased in busulfan-treated mice. Data are expressed as mean ± SEM (n = 3 each). *P <0.05, **P <0.01, ***P <0.001 vs. control (Welch’s t-test).

### Evaluation of testicular histology

The histopathological analysis of the seminiferous tubules showed that busulfan-treated mice accounted for approximately half of seminiferous tubules with spermatogenic cell loss; in contrast, this was not observed in control mice (Fig. 3A, B). In busulfan-treated mice, testicular toxicity was characterized by the vacuolization of Sertoli cells and degeneration of spermatocytes and spermatids (Fig. 3B). PAS-positive effusions were also observed in the interstitium of the seminiferous tubules of these mice (Fig. 3B).

**Figure 3.**
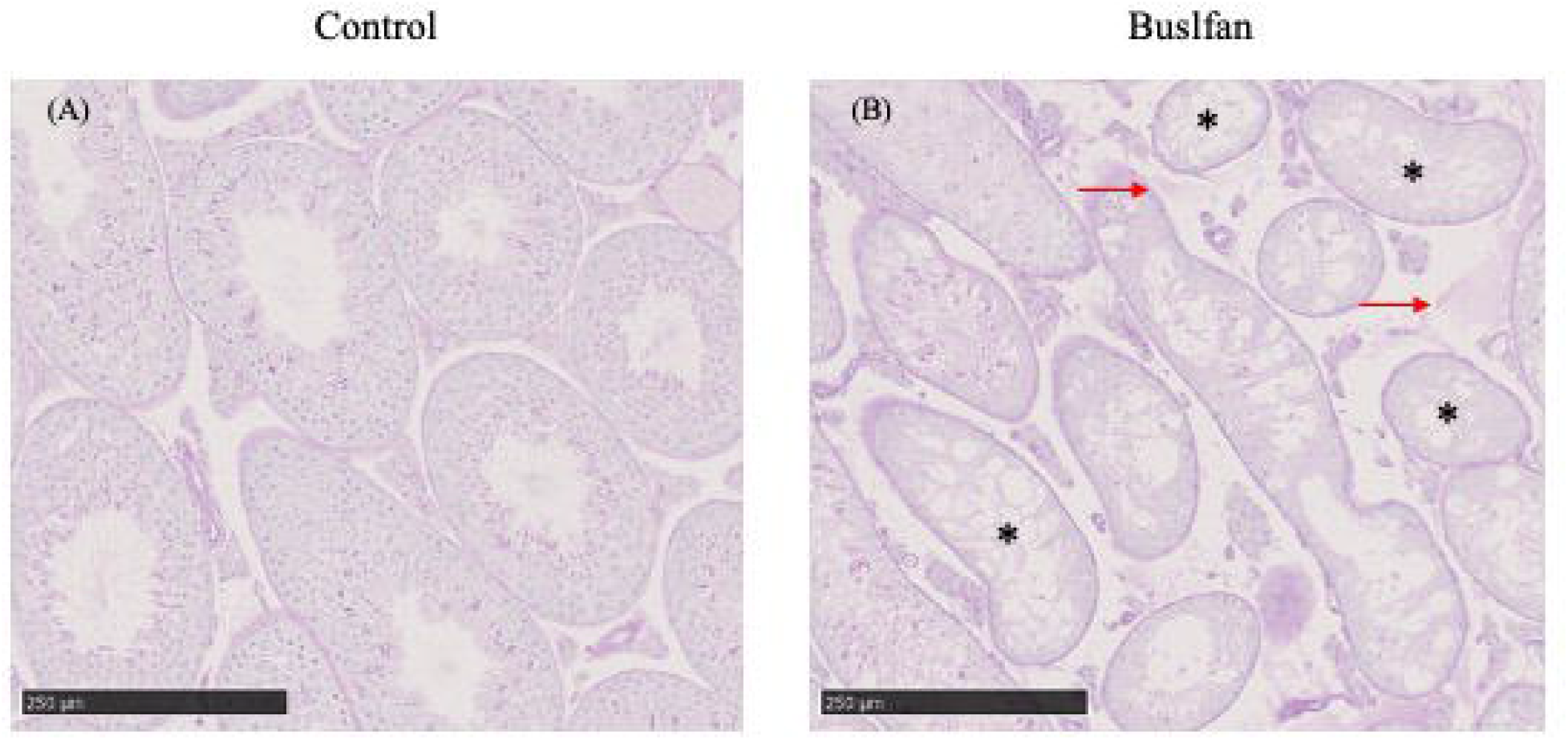
Photomicrographs of mouse testes cross-sections. The tissues were stained with periodic acid–Schiff–hematoxylin (PAS-H) and analyzed at 200 × magnification (scale bar = 250 µm). (A) Photomicrographs of cross-sections from control mice. (B) Photomicrographs of cross-sections from busulfan-treated mice. The samples were collected 28 days post-treatment. Most busulfan-treated mice showed depletion of spermatogenic cells and severe disruption of seminiferous tubular structures in the testes; however, approximately 40% of these structures contained only somatic cells. PAS-positive effusions were observed in the interstitium of busulfan-treated mouse testes (arrows in red, n = 3 each). *shows spermatogenic cell loss in the seminiferous tubules.

## Discussion

The histopathology and weight of the testes are the best non-clinical endpoints to detect testicular toxicity in experimental animals. However, testicular toxicity testing has been a challenge due to the lack of simple and robust screening methods; thus, to evaluate non-clinical toxicity in vivo, the number of animals to be used cannot be reduced. The present study demonstrated the significance of developing a non-invasive method for testicular toxicity evaluation using an infertility mouse model; this model made more feasible predictions even with decreased time and resources.

The present study used well-established, busulfan-treated mice to produce a male infertility model. Busulfan, an alkylating agent, can inhibit cell division by binding to one of the DNA strands, making spermatogenic cells with high cell division rates susceptible to busulfan treatment. A single i.p. of busulfan eliminates almost all endogenous spermatogenic cells in mice; this has become the most commonly technique to induce infertility (Nakata et al., 2020, Gutierrez et al., 2016, Wang et al., 2010). To sterilize the mice, a single-dose i.p. of busulfan should exceed 30 mg/kg (Nakata et al., 2020, Gutierrez et al., 2016, Wang et al., 2010); however, this notably increases the mortality rate of mice (Wang et al., 2010). Previous studies used 50 % DMSO in saline or water as a vehicle control to dissolve busulfan (Nakata et al., 2020, Gutierrez et al., 2016, Wang et al., 2010); however, this cannot be applied to toxicity studies. In the present study, we established a method to dissolve busulfan in 10% DMSO and 90% saline solution. As previously described, to deplete spermatogenic cells without lethality, we administered two i.p. of busulfan at 3-hr intervals (1 mg/mL busulfan × 20 mL/kg per injection) and a total dose of 40 mg/kg body weight (Xie et al., 2020); this did not have a lethal effect on a total of 36 mice, even in the preliminary test (data not shown).

Histopathological analysis is the gold standard for evaluating testicular toxicity (Takahashi & Matsui, 1993). In non-clinical studies, changes in testes weight can be a sensitive indicator of moderate damage to the testes (Lanning et al., 2002). However, this approach requires many animals for time-course toxicity testing, which includes observation of recovery from anatomic lesions over time; this necessitates non-invasive methods of measurement. MRI is often used as a non-invasive alternative to evaluate testicular architecture and pathology. A novel, high-performance compact MRI with a high-field permanent magnet has been utilized in several non-clinical studies, encompassing a wide range of applications in hepato-, nephro-, and neurotoxicity studies, among others (Taketa et al., 2015, Tempel-Brami et al., 2015). However, no reports have been made about non-invasive 3D images of the testes from in vivo MRI analysis. To the best of our knowledge, this is the first study to demonstrate the utility of MRI analysis in evaluating testicular toxicity in a mouse model of busulfan-induced spermatogenic disorder; busulfan causes extensive degeneration of seminiferous tubules (de Rooij & Kramer, 1970, Bucci & Meistrich, 1987).

In the present study, we obtained 3D images in vivo by using a T1-weighted, 3D gradient-echo MRI sequence. This gapless sequence has several advantages over the standard multi-slice imaging procedure. Gapless image acquisitions are most profitable in grasping small organs, such as mouse testes, because a slice gap is inevitable in the standard multi-slice imaging procedure. Custom image analysis scripts were created for the automated segmentation of the testes in situ. The segments were serially measured, and the total testicular volume was calculated by reconstructing these areas. The T1-weighted, 3D gradient-echo sequence supported our observations on the gross pathology of the testes: the total testicular volume in busulfan-treated mice was significantly smaller than in control mice. The seminiferous tubule volume matches the testicular volume because the ratio of the total testicular volume to the total seminiferous tubule volume is similar between individuals irrespective of testes size (Nakata et al., 2015). Histopathological analysis revealed that the significant reduction in testes weight among busulfan-treated mice was mostly due to spermatogenic cell depletion in the seminiferous tubules. On the other hand, imaging provides 3D quantitative data on tissue changes that cannot be easily acquired using traditional pathological approaches. In vivo 3D imaging of the testes can help predict the effects of pharmaceuticals, such as anti-cancer drugs, on testicular histopathology, even without dissection. In the future, MRI systems can provide useful temporal information to evaluate testicular toxicity in the same animals, potentially reducing the number of animals used in toxicological studies.

Finally, compared to that of control mice, the signal on T1-weighted images significantly increased in the testes of busulfan-treated mice. Several clinical studies have shown that high T1 signals (T1 shortening) are detected in high concentrations of macromolecules, such as in mucin-containing tumors (Gaeta et al., 2002). The oligosaccharides attached to the peptide chains within the thick mucin causes T1 shortening either by themselves or through water–macromolecule interactions, which can increase T1 signal intensity within tumors (Wei et al., 2022). Histopathological analysis revealed PAS-positive effusions in the interstitium of the testes of busulfan-treated mice. Given the principle of PAS staining, these effusions may be composed of glycoprotein-like substances. Therefore, the results of the staining may be related to T1 shortening due to a high concentration of glycoproteinaceous content.

In conclusion, the present study demonstrated that small animal imaging in vivo can detect morphological and anatomical changes in the testes, using an infertility mouse model to better understand non-clinical testicular toxicity. However, the study had limitations. The data obtained was insufficient for a time-course evaluation. Spermatogenesis is a continuous, cyclical, and synchronized process that occurs in the epithelium of the seminiferous tubules of the testes, spanning approximately 35 days in mice and 74 days in humans (Fayomi & Orwig, 2018). Chemicals may influence the testes at any point in the cycle. Therefore, it is difficult to predict when and where testicular toxicity may occur. Testicular toxicity should not be evaluated at only one time point but at many time points. Further investigations are needed to evaluate sequential non-invasive 3D imaging by MRI analysis. This will enable researchers to examine the longitudinal progression of the signs of testicular toxicity and its reversibility in the same animals treated with chemicals.

## Acknowledgment

The authors thank Mr. Yoshiharu Tsuru, Mr. Itaru Higuchi, and Mr. Takahiro Aoki (Primetech Co. Ltd., Tokyo, Japan) for their technical assistance. We thank Editage (https://www.editage.jp) for their English language editing service. This study was supported by the Japan Agency for Medical Research and Development (grant number 21mk0101210j0001 to Satoshi Yokota).

## Authors’ contributions

SY conceived and designed the experiments. SK supervised the study. SY, HM, and KS performed the experiments. SY, TH, and TW discussed the results of the histological and MRI data. SY drafted the manuscript. TH, TW, YT, and SK critically revised the manuscript for intellectual content.

## References

Bucci, L.R. and Meistrich, M.L. (1987): Effects of busulfan on murine spermatogenesis: cytotoxicity, sterility, sperm abnormalities, and dominant lethal mutations. Mutat Res, 176, 259–268.

Daneshgaran, G., Lo, A.Y., Paik, C.B., Cooper, M.N., Sung, C., Jiao, W., Park, S.Y., Ni, P., Yu, R.P., Vorobyova, I., Jashashvili, T., Hong, Y.K., Kim, G.H., Conti, P.S., Chai, Y. and Wong, A.K. (2019): A Pre-clinical Animal Model of Secondary Head and Neck Lymphedema. Scientific reports, 9, 18264.

de Rooij, D.G. and Kramer, M.F. (1970): The effect of three alkylating agents on the seminiferous epithelium of rodents. I. Depletory effect. Virchows Arch B Cell Pathol, 4, 267–275.

Fayomi, A.P. and Orwig, K.E. (2018): Spermatogonial stem cells and spermatogenesis in mice, monkeys and men. Stem Cell Res, 29, 207–214.

Gaeta, M., Vinci, S., Minutoli, F., Mazziotti, S., Ascenti, G., Salamone, I., Lamberto, S. and Blandino, A. (2002): CT and MRI findings of mucin-containing tumors and pseudotumors of the thorax: pictorial review. Eur Radiol, 12, 181–189.

Gutierrez, K., Glanzner, W.G., Chemeris, R.O., Rigo, M.L., Comim, F.V., Bordignon, V. and Gonçalves, P.B. (2016): Gonadotoxic effects of busulfan in two strains of mice. Reprod Toxicol, 59, 31–39.

Lanning, L.L., Creasy, D.M., Chapin, R.E., Mann, P.C., Barlow, N.J., Regan, K.S. and Goodman, D.G. (2002): Recommended approaches for the evaluation of testicular and epididymal toxicity. Toxicologic pathology, 30, 507–520.

Nakata, H., Nakano, T., Iseki, S. and Mizokami, A. (2020): Three-Dimensional Analysis of Busulfan-Induced Spermatogenesis Disorder in Mice. Front Cell Dev Biol, 8, 609278.

Nakata, H., Wakayama, T., Sonomura, T., Honma, S., Hatta, T. and Iseki, S. (2015): Three-dimensional structure of seminiferous tubules in the adult mouse. J Anat, 227, 686–694.

Natt, O., Watanabe, T., Boretius, S., Radulovic, J., Frahm, J. and Michaelis, T. (2002): High-resolution 3D MRI of mouse brain reveals small cerebral structures in vivo. J Neurosci Methods, 120, 203–209.

Pichler, B.J., Kolb, A., Nägele, T. and Schlemmer, H.P. (2010): PET/MRI: paving the way for the next generation of clinical multimodality imaging applications. J Nucl Med, 51, 333–336.

Pittala, S., Krelin, Y. and Shoshan-Barmatz, V. (2018): Targeting Liver Cancer and Associated Pathologies in Mice with a Mitochondrial VDAC1-Based Peptide. Neoplasia, 20, 594–609.

Scialli, A.R., Clark, R.V. and Chapin, R.E. (2018): Predictivity of Nonclinical Male Reproductive Findings for Human Effects. Birth Defects Res, 110, 17–26.

Simmons, G.H., Christenson, J.M., Kereiakes, J.G. and Bahr, G.K. (1975): A non-linear programming method for optimizing parallel-hole collimator design. Phys Med Biol, 20, 771–788.

Sotos, J.F. and Tokar, N.J. (2012): Testicular volumes revisited: A proposal for a simple clinical method that can closely match the volumes obtained by ultrasound and its clinical application. Int J Pediatr Endocrinol, 2012, 17.

Takahashi, M. and Matsui, H. (1993): Mechanisms of testicular toxicity. J Toxicol Pathol, 6, 161–174.

Taketa, Y., Shiotani, M., Tsuru, Y., Kotani, S., Osada, Y., Fukushima, T., Inomata, A. and Hosokawa, S. (2015): Application of a compact magnetic resonance imaging system for toxicologic pathology: evaluation of lithium-pilocarpine-induced rat brain lesions. J Toxicol Pathol, 28, 217–224.

Tempel-Brami, C., Schiffenbauer, Y.S., Nyska, A., Ezov, N., Spector, I., Abramovitch, R. and Maronpot, R.R. (2015): Practical Applications of in Vivo and ex Vivo MRI in Toxicologic Pathology Using a Novel High-performance Compact MRI System. Toxicologic pathology, 43, 633–650.

Wang, D.Z., Zhou, X.H., Yuan, Y.L. and Zheng, X.M. (2010): Optimal dose of busulfan for depleting testicular germ cells of recipient mice before spermatogonial transplantation. Asian J Androl, 12, 263–270.

Wei, P.K., Gupta, M., Tsai, L.L., Lee, K.S., Jaramillo, A.M., Smith, M.P., LeGout, J.D. and Shenoy-Bhangle, A.S. (2022): Spectrum of MRI Features of Mucin-producing Neoplasms in the Abdomen and Pelvis. Radiographics, 42, 469–486.

Xie, Y., Deng, C.C., Ouyang, B., Lv, L.Y., Yao, J.H., Zhang, C., Chen, H.C., Li, X.Y., Sun, X.Z., Deng, C.H. and Liu, G.H. (2020): Establishing a nonlethal and efficient mouse model of male gonadotoxicity by intraperitoneal busulfan injection. Asian J Androl, 22, 184–191.

Zhang, J., Yarowsky, P., Gordon, M.N., Di Carlo, G., Munireddy, S., van Zijl, P.C. and Mori, S. (2004): Detection of amyloid plaques in mouse models of Alzheimer’s disease by magnetic resonance imaging. Magn Reson Med, 51, 452–457.

